# A Reaction-Diffusion Network model predicts a dual role of Cactus/IκB to regulate Dorsal/NFκB nuclear translocation in *Drosophila*

**DOI:** 10.1101/2020.01.18.909416

**Authors:** C.D.T. Barros, M.A. Cardoso, P.M. Bisch, H.M. Araujo, F.J.P. Lopes

**Affiliations:** Laboratório Nacional de Computação Científica (LNCC), Petrópolis, Brasil.; Laboratório de Física Biológica, Instituto de Biofísica Carlos Chagas Filho (IBCCF), Universidade Federal do Rio de Janeiro (UFRJ) - Rio de Janeiro, Brasil; Laboratório de Biologia Molecular do Desenvolvimento, Instituto de Ciências Biomédicas, Universidade Federal do Rio de Janeiro (UFRJ), Centro de Ciências da Saúde - Rio de Janeiro, Brasil; Grupo de Biologia do Desenvolvimento e Sistemas Dinâmicos, Campus Duque de Caxias Professor Geraldo Cidade, Universidade Federal do Rio de Janeiro (UFRJ) – Duque de Caxias, Brasil

**Keywords:** NFkappaB, Dorsal gradient, morphogen, mathematical model, Toll

## Abstract

Dorsal-ventral patterning of the *Drosophila* embryo depends on the NFκB superfamily transcription factor Dorsal (Dl). Toll receptor activation signals for degradation of the IκB inhibitor Cactus (Cact), leading to a ventral-to-dorsal nuclear Dl gradient. Cact is critical for Dl nuclear import, as it binds to and prevents Dl from entering the nuclei. Quantitative analysis of *cact* mutants revealed an additional Cact function to promote Dl nuclear translocation in ventral regions of the embryo. To investigate this dual Cact role, we developed a predictive model based on a reaction-diffusion regulatory network. This network considers non-uniform Toll activation as well as Toll-dependent Dl nuclear import and Cact degradation. In addition, it incorporates translational control of Cact levels by Dl, a Toll-independent pathway for Cact regulation and reversible nuclear-cytoplasmic Dl flow. Our model successfully reproduces wild-type data and emulates the Dl nuclear gradient in mutant *dl* and *cact* allelic combinations. Our results indicate that the dual role of Cact depends on targeting distinct Dl complexes along the dorsal-ventral axis: In the absence of Toll activation, free Dl-Cact trimers inhibit direct Dl nuclear entry; upon ventral-lateral Toll activation, Dl-Cact trimers are recruited into predominant signaling complexes and promote active Dl nuclear translocation. Simulations suggest that Toll-independent regulatory mechanisms that target Cact are fundamental to reproduce the full assortment of Cact effects. Considering the high evolutionary conservation of these pathways, our analysis should contribute to understand NFκB/c-Rel activation in other contexts such as in the vertebrate immune system and disease.

## INTRODUCTION

In a developing organism, tissues are often patterned by long-range and spatially graded signaling factors called morphogens, which carry the positional information necessary to control gene expression. Morphogens act in a concentration-dependent manner to activate or repress target genes (Driever and Nusslein-Volhard, 1988; Lopes *et al*., 2008; Roth *et al*., 1991; Struhl *et al*., 1989). Therefore, precisely defining the amount of activated morphogen is crucial to determining their effects. In the *Drosophila* syncytial blastoderm, dorsal-ventral (DV) patterning depends on the nuclear localization gradient of Dorsal (Dl), a NFκB superfamily transcription factor homologous to mammalian c-Rel. Dl acts in a concentration-dependent manner to activate or repress target genes, defining three main territories of the embryo DV axis: the ventral mesoderm, lateral neuroectoderm, and dorsal ectoderm (Rushlow and Shvartsman, 2012). A ventral-to-dorsal activity gradient of the Toll cell surface receptor provides the activating signal for DV patterning. Dl nuclear translocation is controlled by Cactus (Cact), a cytoplasmic protein related to mammalian IκB. In the absence of Toll signals, Cact binds to Dl and impairs its nuclear translocation (Bergmann *et al*., 1996). Activated Toll receptors on the ventral and lateral regions of the embryo lead to Cact phosphorylation followed by ubiquitination and degradation, resulting in dissociation of the Dl-Cact complex and Dl nuclear import. Different Dl levels subdivide the embryonic DV axis in target gene expression domains, defined by distinct thresholds of sensitivity to control by Dl (Stein and Stevens, 2014). Thus, the amount of Dl in the nuclei is key to defining ventral, lateral and dorsal territories of the embryo.

In attempts to explain the complex Dl regulatory signaling network, mathematical models were developed to simulate the Dl nuclear gradient. Early models simulated gradient profiles throughout nuclear division cycles 10 to 14 in wild-type embryos, with parameters constrained by experimental data from endogenous nuclear Dl (nDl) levels in fixed embryos or live imaging of Dl-GFP (Kanodia *et al*., 2009; Liberman *et al*., 2009). Their results showed that the nuclear Dl gradient is dynamic, increasing in amplitude from cycle 10 to cycle 14, without significantly changing its shape. Subsequently, it was shown that both the nDl gradient amplitude and basal levels oscillate throughout early embryonic development (Reeves *et al*., 2012). Recently, it was proposed that facilitated diffusion along the DV axis, or “shuttling” via Dl-Cact complexes, plays a role in nDl gradient formation (Carrell *et al*., 2017). Therefore, these analyses suggest that the establishment of the nDl gradient is a complex process yet to be fully understood.

Former initiatives modeling the nDl gradient focused predominantly on processes regulated by Toll. However, several proteins that impact Dl nuclear localization act independent of Toll-receptor activation. For instance, adaptor proteins that are required to transduce Toll signals, such as Tube, Pelle and Myd88, form pre-signaling complexes that depend on the interaction with cytoskeletal elements (Edwards *et al*., 1997; Galindo *et al*., 1995; Towb *et al*., 1998) and the membrane (Ji *et al*., 2014; Marek and Kagan, 2012). In addition, alterations in Cact concentration and Calpain A protease activity perturb the nDl gradient (Fontenele *et al*., 2013; Roth *et al*., 1991). Dl itself can flow into and out of the nucleus in the absence of Toll (Carrell *et al*., 2017; DeLotto *et al*., 2007), with the potential to contribute significantly to nDl concentration. Since Cact plays a critical role on the control of Dl nuclear translocation, understanding how Cact is regulated is paramount.

Two pathways have been reported to regulate Cact levels: the Toll dependent pathway, considered in all the models to date, that leads to Cact N-terminal phosphorylation and degradation through the proteasome, and a Toll-independent pathway that targets Cact C-terminal sequences for phosphorylation (Belvin *et al*., 1995). This originally termed “signal independent pathway for Cact degradation” (Belvin *et al*., 1995; Bergmann *et al*., 1996; Liu *et al*., 1997; Reach *et al*., 1996) acts in parallel to Toll-induced signaling (Moussian and Roth, 2005). However, several reports indicate that the signal-independent pathway is regulated by Casein kinase II (Liu *et al*., 1997; Packman *et al*., 1997), the BMP protein encoded by *decapentaplegic* (Araujo and Bier, 2000; Carneiro *et al*., 2006) and by the calcium-dependent modulatory protease Calpain A (Fontenele *et al*., 2009). Moreover, Calpain A generates a Cact fragment that is more stable than full-length Cact (Araujo *et al*., 2018; Fontenele *et al*., 2013), indicating that the Toll-independent pathway may perform unexplored roles in nDl gradient formation.

Recently, by analyzing the effects of a series of *cact* and *dorsal (dl)* loss-of-function alleles, we detected a novel function of Cact to promote Dl nuclear translocation. In addition to inhibiting Dl nuclear import, Cact also acts to favor Dl nuclear translocation where Toll signals are high (Cardoso *et al*., 2017). However, the mechanism behind this effect is unclear. Since previous mathematical models for Dl gradient formation did not encompass the two mechanisms that regulate Cact function, here we propose a new reaction-diffusion model to describe this signaling network. We take into account non-uniform activation of Toll as in current models, two Dl nuclear translocation mechanisms, translational regulation of Cact by Dl (Govind *et al*., 1993; Kubota and Gay, 1995) and the two pathways leading to Cact regulation: the Toll-regulated and Toll-independent pathways. Using a Genetic Algorithm and experimental data from wild-type *Drosophila* embryos, we calibrate the model parameters, such as kinetic constants, diffusion coefficients, Toll activation profile and total Dorsal concentration in the embryo. The optimized parameters are then used to reproduce and understand the nDl patterns of single and double mutants for *cactus* and *dorsal* genes. Our model analysis indicates that, in the ventral region, Cact favors Dl nuclear translocation by the formation of Toll-responsive Dl-Cact complexes that signal to Dl nuclear localization, while throughout the entire embryo Cact binds to and prevents free Dl from entering the nuclei.

## RESULTS

### A reaction-diffusion model that discriminates two routes for Dorsal nuclear localization highlights the distinctive contribution of Toll-activated versus free Dl flow to the nDl gradient

Embryonically translated from maternal mRNAs deposited in the egg, Dl protein enters the nucleus through two different mechanisms: actively imported upon degradation of the Cact inhibitor by Toll signals, and by direct flow in and out of the nucleus (DeLotto *et al*., 2007). Immunological detection of Dl in fixed embryos, or visualization of fluorescently tagged Dl in live tissue, does not discriminate the Dl dimers that enter nuclei in response to Toll versus those that enter by direct flow. In order to investigate how these different Dl nuclear entry modes impact the nuclear Dl (nDl) gradient we built a model that discriminates these two mechanisms. In the first mechanism described, activated Toll recruits a 2Dl-Cact trimeric complex (DlC) to the Tl-signaling complex (DlCT) (represented by the reversible reaction related to kinetic constants k7 and k8, Fig. 1), leading to the irreversible dissociation of phosphorylated Dl from Cact (k9), Cact degradation (k10), and consequent entry of a Dl dimer in the nucleus (k11). In a final step of this route the Dl dimer returns to the cytoplasm (k12). The second nuclear entry mode depends on the direct, reversible flow of free Dl dimers in and out of the nucleus (k3 and k4). In the cytoplasm free Dl dimers can associate to Cact (k5 and k6), generating DlC that keeps Dl form entering the nuclei. In addition, the model includes Dorsal-mediated translational regulation of Cactus (k1), as well as Toll-independent Cact processing that may generate Cact fragments with different activities and/or leads to Cact degradation (k2) (Belvin *et al*., 1995; Fontenele *et al*., 2013). This model reproduces the characteristic ventral-to-dorsal nDl gradient displayed in wild-type embryos during cycle 14 (Fig. 2A-C; Table 1). Furthermore, it allows discriminating the contribution of each Dl nuclear transport mechanism along the entire DV axis to generate the compounded nDl pattern (Fig. 2D). Thus, a homogenous direct flow of Dl dimers into the nucleus is observed along the DV axis (Fig. 2D, nDl^0^), while Toll-induced nuclear Dl displays ventral-to-dorsal asymmetry (Fig. 2D, nDl*). Consequently, in the ventrolateral region the contribution from the Toll-induced mechanism to nDl levels is much larger than from direct flow, resulting in a significant gradient of nuclear Dorsal. On the other hand, in the dorsal region the contribution from direct flow is predominant.

**Figure 1:**
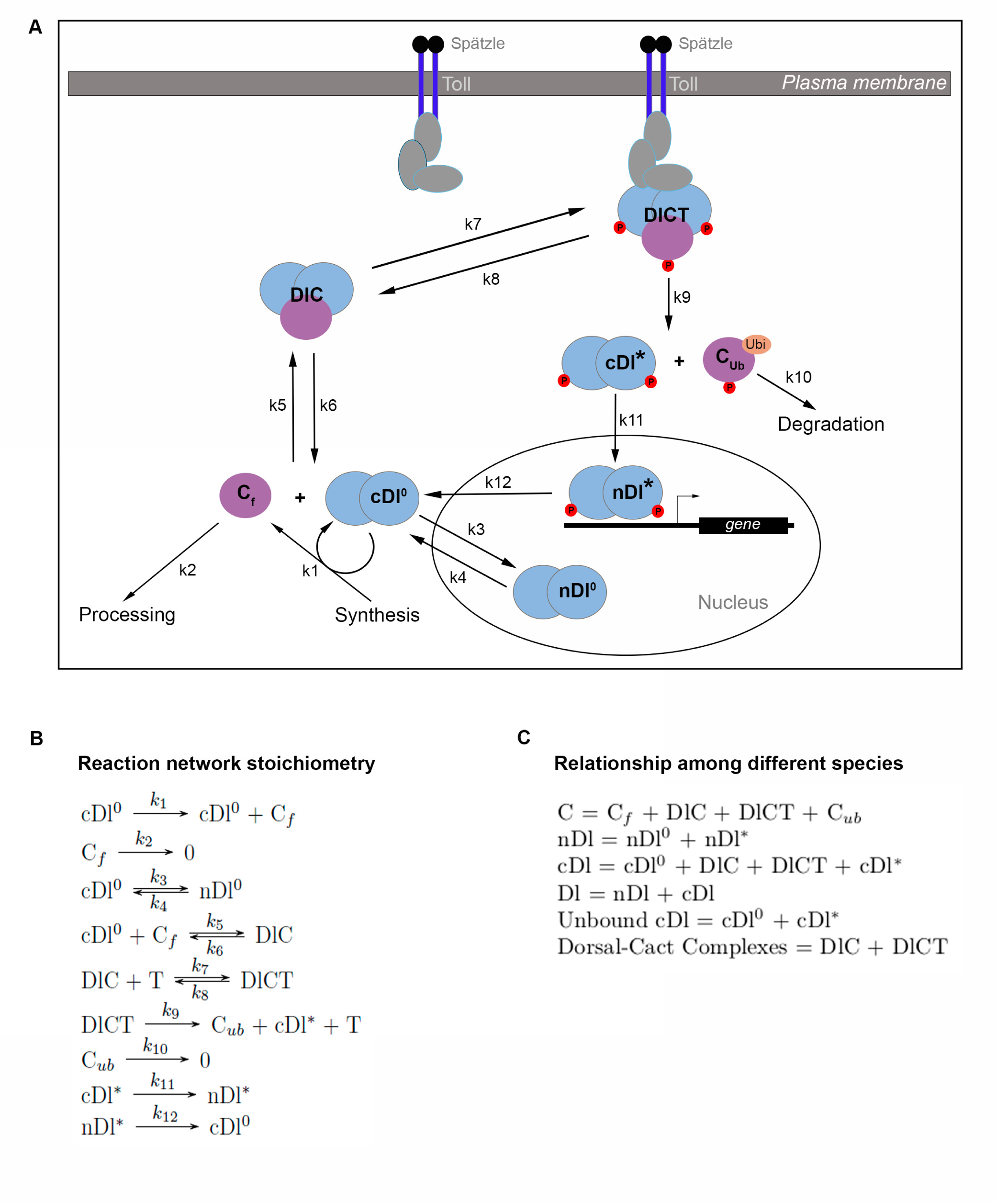
Mathematical modeling of nuclear Dorsal gradient in *Drosophila* embryos. (A) Schematic representation of the Reaction-Diffusion Network Model regulating nuclear Dorsal (nDl) localization. Kinetic constants k1 and k2 mediate synthesis and degradation of the inhibitor Cactus, respectively. Dl dimers can enter and leave the nucleus by direct flow, independently of Toll (k3 and k4). k5 and k6 mediate reversible binding between cytoplasmic Dl dimers (cDl^0^) and free Cactus (C_f_) to form trimeric complexes (DlC). DlCT complex is reversibly formed by interaction between DlC and activated Toll membrane receptor (k7 and k8). Toll activation induces Dl and Cactus phosphorylation, releasing their complexes. This irreversible reaction is controlled by k9. Cytoplasmic phosphorylated Dl dimers (cDl*) enter the nucleus (nDl*) while phosphorylated and ubiquitinated Cactus (C_ub_) is degraded by the proteasome (k10). k12 controls nDl* output from the nucleus. T represents activated Toll receptor. (B) Detailed reaction network stoichiometry. (C) Key relationships among model species. Total Cactus (C) is the sum of all species that contain Cactus. Total nuclear Dorsal (nDl) is the sum of nDl^0^ and nDl*. Total cytoplasmic Dorsal (cDl) includes free cDl, cDl*, plus the two DlC and DlCT complexes. Total Dorsal is the sum of nDl and cDl. Model was solved for cleavage cycle 14 embryos.

**Figure 2:**
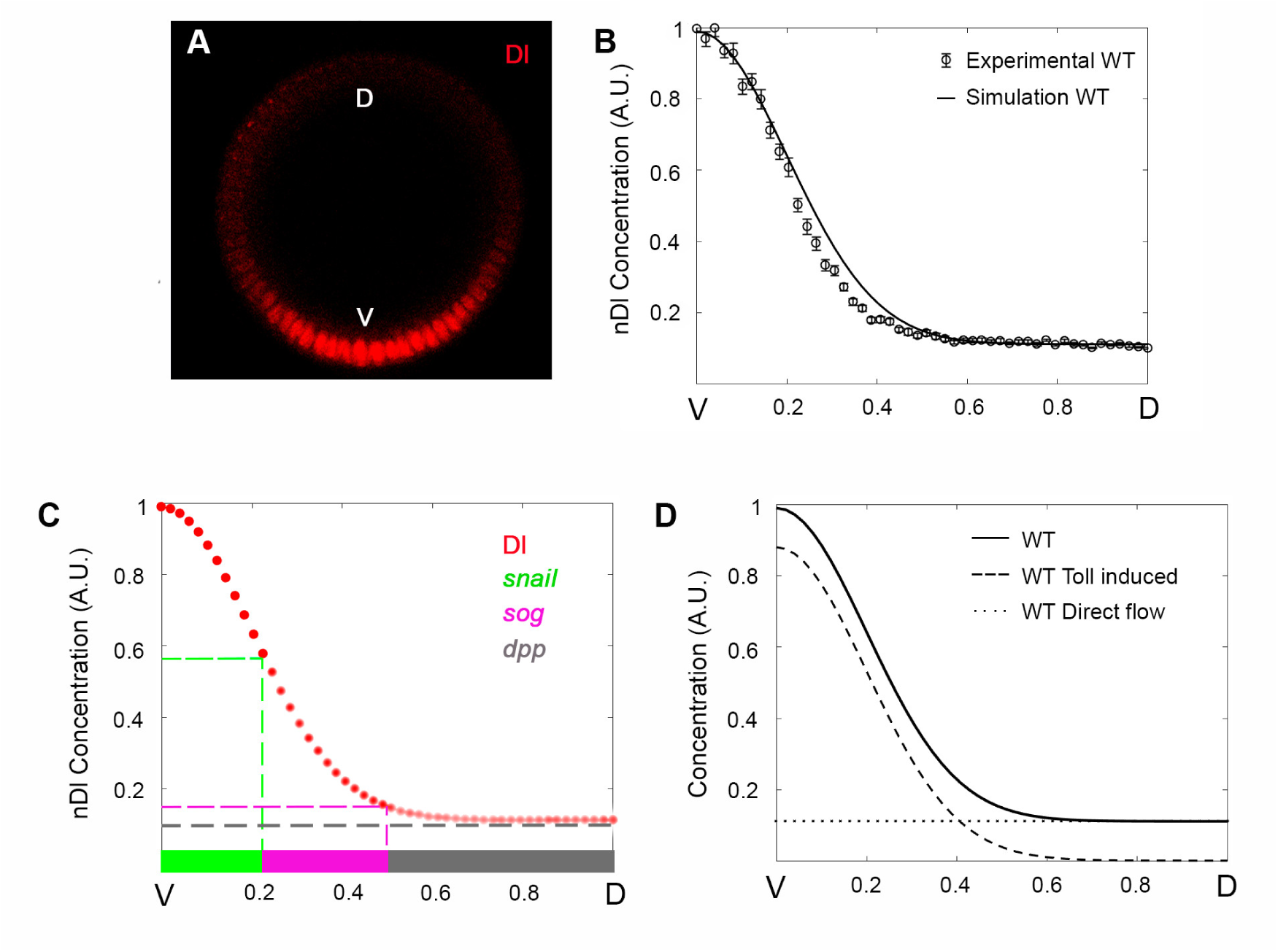
The Reaction-Diffusion Network Model reproduces the wild type nDl gradient profile and discriminates two different Dl nuclear entry modes. (A) Optical section of a wild type (WT) embryo stained for Dorsal protein. (B) Nuclear Dl fluorescence intensity was extracted from sections as in A, measured and plotted as half gradients (circle). The black curve displays model simulation. The y axis represents nDl fluorescence intensity along the ventral-to-dorsal (V-D) embryonic axis (x axis). Data are mean±s.e.m. (C) High to low nDl levels (red circles) define different DV territories: ventral mesoderm represented by *snail* expression (green), lateral neuroectoderm represented by *short gastrulation* (*sog*, magenta) and dorsal ectoderm defined by *decapentaplegic* expression (*dpp*, grey). (D) Simulations discriminate nuclear Dl that enters the nucleus by direct flow (nDl^0^, dotted curve) or induced by Toll (nDl*, dashed curve). Black curve indicates total nDl model simulation, as in B. Ventral (V) region to the left, dorsal (D) to the right.

**Table 1:**
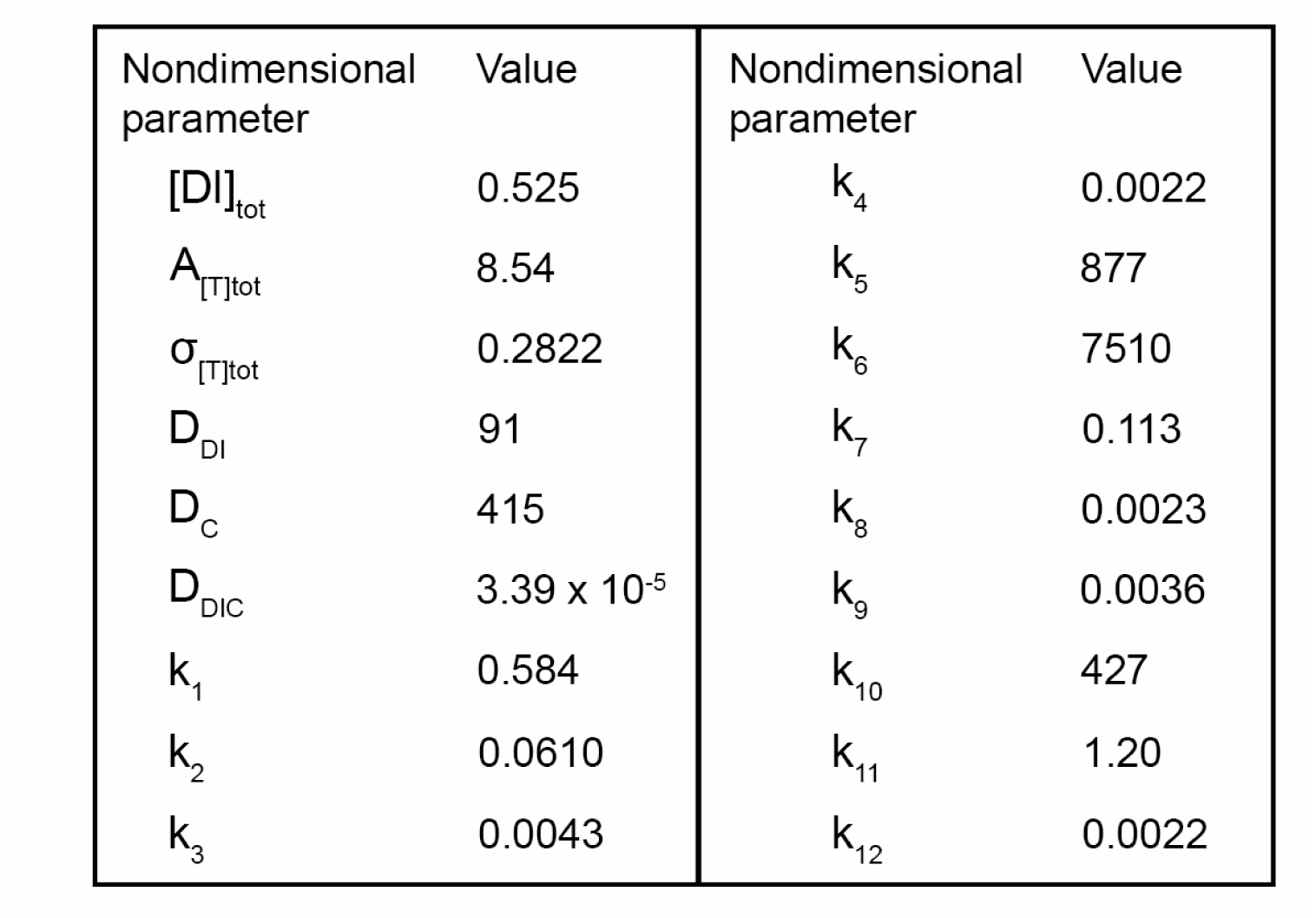
Dimensionless model parameters. Model parameters calibrated to simulate wild-type data. The dimensionless concentrations were obtained by normalizing the experimental data, as described in Methods. Specifically, for Dl concentration and peak Toll concentration, normalization is obtained by dividing these values by the experimental nuclear Dl concentration in the most ventral region. For quantities involving length, including diffusion coefficients and the width of the Gaussian that represents the Toll activation profile, the total size of the half-embryo was considered to be unitary length (therefore, each compartment is 1/50 in size). For the quantities involving time, such as diffusion coefficients and kinetic constants, the characteristic time was T = 1.

| Nondimensional parameter | Value | Nondimensional parameter | Value |
| --- | --- | --- | --- |
| $[DI]_{tot}$ | 0.525 | $k_4$ | 0.0022 |
| $A_{[T]tot}$ | 8.54 | $k_5$ | 877 |
| $\sigma_{[T]tot}$ | 0.2822 | $k_6$ | 7510 |
| $D_{DI}$ | 91 | $k_7$ | 0.113 |
| $D_C$ | 415 | $k_8$ | 0.0023 |
| $D_{DIC}$ | $3.39 \times 10^{-5}$ | $k_9$ | 0.0036 |
| $k_1$ | 0.584 | $k_{10}$ | 427 |
| $k_2$ | 0.0610 | $k_{11}$ | 1.20 |
| $k_3$ | 0.0043 | $k_{12}$ | 0.0022 |

One fundamental characteristic of the embryonic Dl gradient is that it is robust to variations in Dl and Cact levels, tolerating a Dl/Cact ratio of 0.3 to 2.0 without a significant effect on embryo viability (Govind *et al*., 1993). In order to test how the amount of total Dl protein impacts its nuclear localization, we simulated nDl levels in embryos generated by mothers heterozygous for a loss-of-function *dl* allele (*dl*[6], Cardoso *et al*., 2017). These embryos (from *dl*[6]/+ mothers) produce less Dl protein than embryos generated from wild-type mothers (Cardoso *et al*., 2017). Even though embryos from *dl*[6]/+ mothers show a reduction in peak nDl in the ventral domain, they do not show a significant change in nDl levels in either the lateral or dorsal regions (Fig. 3A, Cardoso *et al*., 2017; Carrell *et al*., 2017). Our model reproduces this pattern and indicates that different Dl components behave distinctively in this context: by lowering the total amount of Dl (Table 2), Toll-responsive nDl* decreases, while nDl^0^ that enters the nucleus by direct flow does not change significantly compared to wild type (Fig. 3B). Accordingly, when the model is tested for greater reductions of maternal Dl (Fig. 3D) nDl reduces significantly, particularly in the ventral region where nDl* is predominant. Furthermore, the model predicts that the ventral peak of DlCT reduces in this mutant (Fig. S1F), while DlC that is uniform along the DV axis, reduces evenly as compared to wild type (Fig. 3C). This pattern of DlCT and DlC reduction conforms to the fact that *dl* controls Cact levels (represented by k1): once Dl levels drop, so do Cact levels and the amount of both DlCT and DlC. The reduction in DlCT reduces the concentration of nDl* in the ventral region (where DlCT is predominant). On the other hand, even though the reduction in Dl also decreases the amount of Dl that enters the nucleus by direct flow, the consequent Cact decrease leaves more Dorsal dimers free for nuclear translocation.

**Figure 3:**
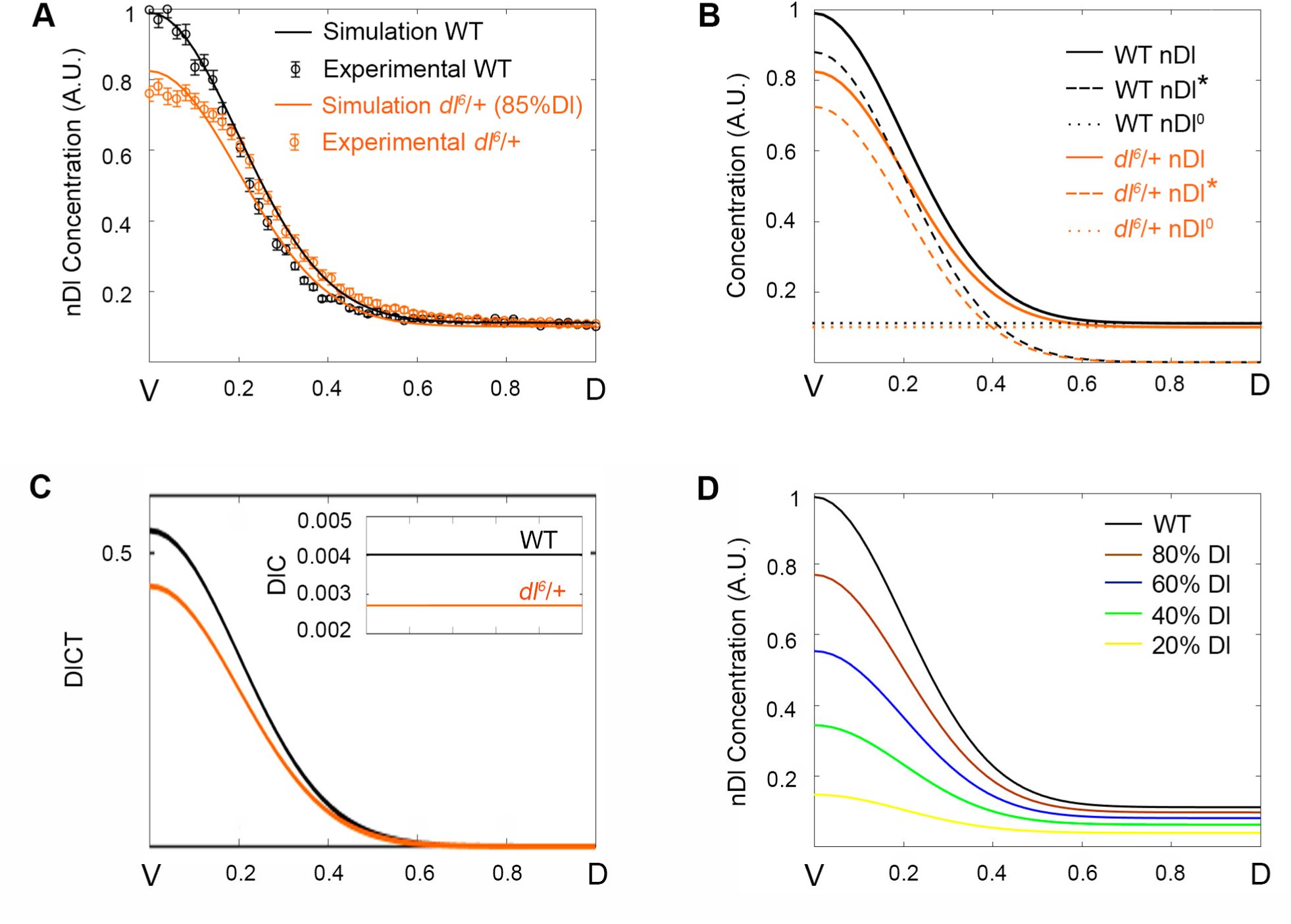
Model reproduces the nuclear Dl gradient in *dl-* mutant embryos. (A) Simulation (solid curves) and experimental data (circle symbols) from mutant (orange) and wild type (black) embryos were plotted in the same graph. Nuclear Dl gradient from *dl^6^*/+ mutant embryos are simulated and fitted using an 85% reduction in total Dorsal protein level. (B) Spatial distribution of Dorsal protein that enters the nucleus by direct flow (nDl^0^, dotted curve), or Toll induced (nDl*, dashed curve) and total nDl. (C) Distribution of DlC and DlCT species amounts for wild type (black) and *dl^6^*/+ (orange) genotypes. (D) nDl gradient simulations resulting from 20% (yellow), 40% (green), 60% (blue), 80% (brown) Dorsal protein reductions compared to a wild type nDl gradient (black).

**Table 2:** Dimensionless parameters for each mutant. Parameters obtained for each mutant simulated, compared to wild type. Only the changeable values are presented, as all other parameters of the model were kept fixed.

| Genotypes | [DI]tot | k1 | [C]tot |
| --- | --- | --- | --- |
| WT | 0.525 | 0.584 | 100% |
| *dl<sup>6</sup>/+* | 0.4605 | 0.584 | 90.70% |
| *cact<sup>A2</sup>/cact<sup>011</sup>* | 0.525 | 0.1825 | 53.40% |
| *dl<sup>6</sup>/cact<sup>A2</sup>* | 0.375 | 0.500 | 71.80% |

### Cact protein acts antagonistically along the DV axis to control the levels of unbound versus complexed Dl

We have previously shown that Cact displays not only a role as inhibitor of Dl nuclear translocation, but also acts to favor nDl localization in ventral nuclei (Fig. 1 in Cardoso *et al*., 2017). In order to use our model to unveil the molecular mechanism behind this dual role exhibited by Cact, we simulated the pattern of hypomorphic *cact* allelic combinations. Initially, we simulated the *cact*[A2]/*cact*[011] genotype, where total Cact levels are reduced in relation to wild type (Table 2), in addition to uncharacterized effects of the mutant alleles (Cardoso *et al*., 2017; Fontenele *et al*., 2013; Roth *et al*., 1991). This genotype shows two opposing effects reproduced by the model: a decrease in nDl in the ventral region and an increase in the lateral and dorsal regions of the embryo (Fig. 4A). The nDl increase in lateral and dorsal regions of the embryo reflects Cact’s classical role described by many authors to inhibit Dl nuclear translocation (Belvin *et al*., 1995; Bergmann *et al*., 1996; Govind *et al*., 1993; Reach *et al*., 1996; Roth *et al*., 1991). Contrarily, the nDl decrease in ventral nuclei, where Toll activation is high, requires a detailed model analysis to be understood.

**Figure 4:**
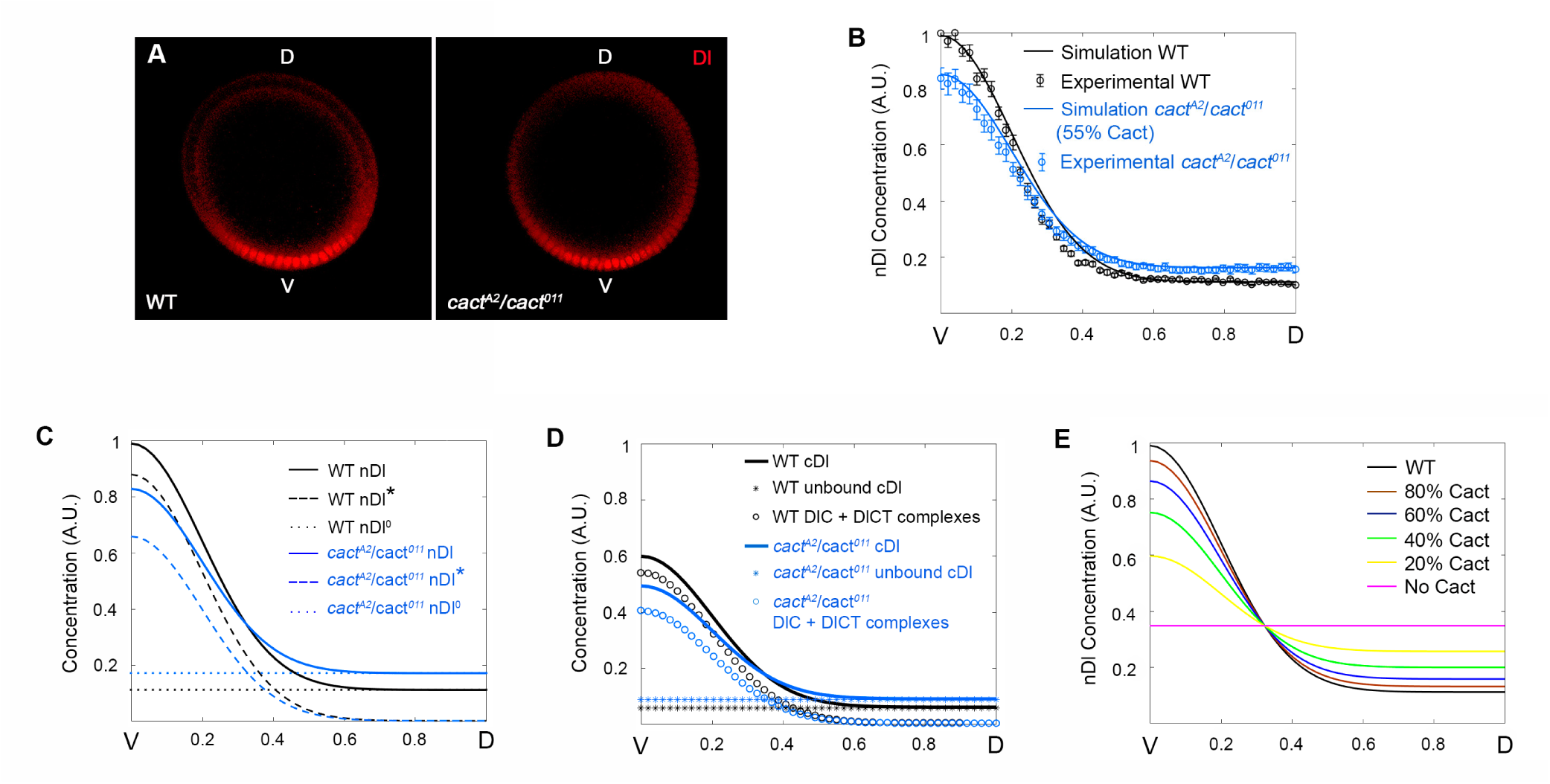
Cact produces distinct effects along the DV axis by controlling unbound versus complexed Dl elements. (A) Simulations (solid curve) and experimental data (circle symbols) from *cactus* mutant (blue) and wild type (black) embryos. Nuclear Dl gradient from *cact^A2^*/*cact^011^* mutant embryos are simulated and fitted using a 55% reduction in Cact protein. (B) Spatial distribution of Dorsal that enters the nucleus by direct flow (nDl^0^, dotted curve), by Toll induced (nDl*, dashed curve) and total nDl. (C) Distribution of Total cDl (defined as cDl^0^ + DlC + DlCT + cDl*; star symbols), sum of the unbound cDl species (cDl^0^ + cDl*) and the DlC + DlCT complexes for wild type (black) and *dl^6^*/+ (orange) genotypes. (D) nDl gradient simulations resulting from 100% (magenta, constant nDl), 20% (yellow), 40% (green), 60% (blue), 80% (brown) Cactus protein reductions are compared to wild type nDl gradient (black).

By analyzing direct and Toll dependent Dl nuclear entry modes separately we found that both are affected, albeit differently, by reducing Cact levels (Fig. 4C). The amount of Dl that directly enters the nuclei (nDl^0^) increases equally along the entire DV axis, consistent with a homogenous increase of free Dl dimers in the cytoplasm (cDl^0^, Fig. S2A). This behavior reflects the well-characterized role of Cact to inhibit Dl nuclear translocation. Conversely, the amount of Dl entering the nucleus in response to the Toll pathway (nDl*) reduces compared to wild type, particularly in the ventral side of the embryo. This nDl* reduction in *cact*[A2]/*cact*[011] reflects loss of a positive role of Cact on Dl nuclear localization. The transition between the positive and negative effects of Cact reduction on Dl nuclear levels is placed exactly at the position along the DV axis where Dl translocation induced by the Toll pathway (dashed blue line in Fig. 4C) matches the amount of Dl that enters by direct flow (dotted blue line in Fig. 4C). This falls inside the lateral domain where intermediate Toll activation takes place. It is important to point out that Dl nuclear import through the Toll pathway is more efficient than via direct flow: the ratio between kinetic constants for Toll-dependent Dl nuclear transport (k11/k12 = 545.45) is higher than via direct flow (k3/k4 = 1.95;Table 1). This is critical to the dual effect observed for *cact*[A2]/*cact*[011). Since the decrease of Dl that translocate to the nucleus in response to Toll (nDl*; Fig. 4C) is greater than the increase in Dl entering the nuclei by direct flow (nDl^0^; Fig. 4C), the resulting effect is a reduction of total nuclear Dl (nDl^0^ + nDl*) in the most ventral region (Fig. 4B-C). Importantly, in addition to the difference in nuclear resident time between nDl^0^ and nDl* suggested by the ratios above, we should note that the effect of nDl^0^ and nDl* on gene expression will also depend on their distinct efficiencies to bind target DNA.

Interestingly, if we consider the different cytoplasmic Dl species that are free (cDl^0^ + cDl*) versus bound to Cact (DlC + DlCT; Fig. 4D), we see that the amount of Cact-bound Dl decreases in the *cact* mutant as compared to wild type. This conforms to experimental data from crude embryonic lysates showing that in *cact* loss-of-function alleles the total amount of Dl-Cact complexes observed in non-reducing gels decreases compared to wild type (Isoda and Nusslein-Volhard, 1994). Accordingly, in our analysis, as the amount of DlCT (Fig. S2F) reduces, a consequent reduction in the amount of modified Dl in the cytoplasm (cDl*) and nuclei (nDl*) is seen (Fig. S2H,I). This nDl* decrease is in agreement with the loss of ventral gene expression (high Dl activated genes) and ventral expansion of lateral gene expression (intermediate Dl level targets) that is observed in this genotype (Cardoso *et al*., 2017).

In order to investigate the effect of more severe reductions in Cact concentration we performed additional simulations with different production rates of Cact, from wild-type rates to zero, that is, in the absence of Cact (Fig. 4E). The model predicts an increase in the Cact dual effect resulting from reductions in Cact levels: In all conditions a more severe loss of Cact lead to less nDl in the ventral side and more nDl in dorsal regions. Furthermore, the model correctly predicts that in Cact absence the gradient is completely lost, reflecting the pattern of an embryo with expanded lateral territories, as shown for some loss-of-function Toll alleles and certain allelic *cact* combinations (Cardoso *et al*., 2017; Ray *et al*., 1991; Anderson *et al*., 1985a; Roth *et al*., 1989). This is also consistent with the main role proposed for Toll being to generate graded Dl activity along the DV axis (Anderson *et al*., 1985a) as opposed to a basic requirement for Dl nuclear translocation.

### *dl/cact* double mutant reduces the levels of Toll-responsive Dorsal-Cactus complexes causing loss of precision in target gene expression domains

Since our results indicate that Cact favors Dl nuclear translocation in the ventral side of the embryo, we decided to challenge this prediction by decreasing Dl and Cact concomitantly. If model prediction is correct, reducing both Cact and Dl should produce a more severe impact in the ventral nDl peak compared to the single *dl*[6]/*+* mutant. This condition is experimentally reached with the *dl*[6]/*cact*[A2] allelic combination, where the levels of both Dl and Cact are reduced (Fig. 5A and S3). We simulated the *dl*[6]/*cact*[A2] mutant by decreasing total Dl (maintained constant throughout the simulation) and the kinetic constant k1, related to Cact production (see Fig. 1A, B; Table 2). The model shows that lowering Cact levels in a *dl*[6]/*+* background produces an additional impact in Dl nuclear translocation in the most ventral region and confirms that Cact has a positive role on Dl nuclear localization. Model discrimination of Toll-induced (nDl*) and direct Dl flow (nDl^0^, Fig. 5B) indicates that in this allelic combination both Dl nuclear translocation mechanisms are reduced.

**Figure 5:**
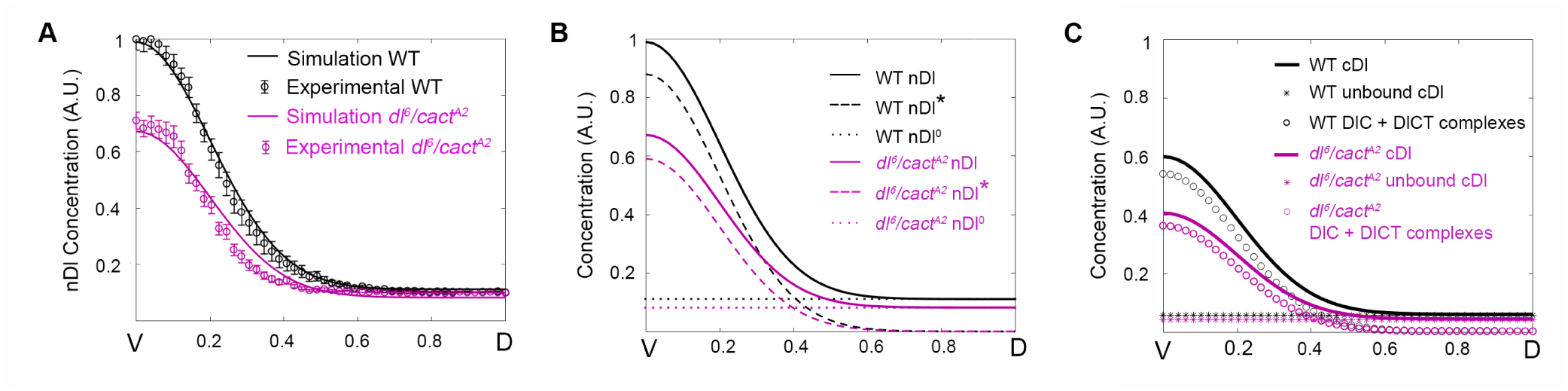
A *dl/cact* double mutant highlights the positive role of Cact on Dl nuclear translocation. (A) Simulations (solid curves) and experimental data (circle symbols) from mutant (purple) and wild type (black) embryos. Nuclear Dl gradients from *dl^6^/cact^A2^* mutant embryos were simulated using simultaneous reduction of 50% Cactus and 70% Dorsal. (B) Spatial distribution of Dorsal that enters the nucleus by direct flow (nDl^0^, dotted curve), by Toll induced (nDl*, dashed curve) and total nDl are shown in wild type (black) and mutant (purple) simulations. (C) Distribution of Total cDl (defined as cDl^0^ + DlC + DlCT + cDl*; star symbols), sum of the unbound cDl species (cDl^0^ + cDl*) and the DlC + DlCT complexes for wild type (black) and *dl^6^/cact^A2^* (purple) genotypes.

In the dorsal region, the effect of reducing Cact production (that has the potential to increase nDl levels, Fig. 4) is almost completely canceled by the reduction in Dl levels (Fig. 5A). Consequently, the dual effect of Cact reduction is no longer observed (compare Fig. 5A and Fig. 4B). By distinguishing the amount of nDl* and nDl^0^, the model shows that nDl resulting from direct flow (nDl^0^) decreases uniformly along the DV axis and that Toll-responsive nDl* decreases ventro-laterally (Fig. 5B). Therefore, in ventral regions the reduction in nDl is compounded by a decrease in both Dl translocation by Toll-dependent and direct flow entry modes. Importantly, the effect on the amount of complexed (DlC + DlCT) versus free cytoplasmic Dl observed in *dl*[6]/*cact*[A2] (Fig. 5C) contrasts from the effect of reducing Cact alone (Fig. 4D). In *dl*[6]/*cact*[A2], both complexed and free Dl are reduced compared to wild type, while in *cact*[A2]/*cact*[011] a reduction in Dl complexes and increase in free Dl is observed. This indicates that a decrease in the relative amount of complexed versus free Dl is critical for the dual effect observed for *cact* loss-of-function.

In considering the role of a morphogen and how the regulatory network impacts morphogen activity it is important to address how the network affects target gene expression domains. A current discussion in the literature concerns whether changes in the slope of the Dl gradient affect target gene expression, especially in the lateral region (Liberman *et al*., 2009; Reeves *et al*., 2012). It is discussed whether differences in the relative amount of nDl in neighboring nuclei are sufficient to define the necessary thresholds for differential gene expression. In wild-type embryos, the *snail/sog* sharp boundary delimits the ventral mesoderm and lateral neuroectoderm domains, positioned at 20% of the DV axis. In *dl*[6]/*cact*[A2] embryos, which lack ventral-lateral boundary precision, the border between the *sog* and *sna* domains is not clearly defined, and *sog* exhibits stochastic expression invading the ventral region (Fig. 6A; Cardoso *et al*., 2017). Therefore, we decided to investigate whether the Dl gradient slope was different among all the genotypes hereby analyzed. To this end, we plotted the derivative of the total nuclear Dorsal concentration (nDl) with respect to the DV axis (Fig. 6B), where the derivative represents the difference in nDl levels between neighboring nuclei along the DV axis. We observed that, although all nDl gradients show similar shape, highest slope values are different for each genotype. Indeed, wild type achieves the greatest absolute slope, whereas the gradient displayed by the *dl*[6]/*cact*[A2] genotype shows the lowest slope (Fig. 6C).This result suggests that in wild-type embryos, the lateral adjacent nuclei exhibit a greater difference in the relative amount of nDl than *dl*[6]/*cact*[A2], which could lead to a precise definition of the embryo’s domains in the control, while the mutant displays less precise domains. This analysis indicates that the relative concentration of nuclear Dl between neighboring nuclei indeed plays a role for domain precision. Conversely, loss of precision in the *sna/sog* border is not observed in *dl*[6]/+ and *cact*[A2]/*cact*[011] embryos (Cardoso *et al*., 2017), indicating that these two mutants still exhibit sufficiently large slopes to define nuclear interdomain differences. Importantly, it has been shown that the refinement of ventral-lateral gene expression borders is also mediated by RNA polymerase II stalling on transcription start sites (Bothma *et al*., 2011; Levine, 2011; Zeitlinger *et al*., 2007), as reported for *short gastrulation (sog)* (Bothma *et al*., 2011). Therefore, a combined effect of distinct nDl levels and transcriptional processes may define the precise gene expression border between adjacent ventral and lateral nuclei of the Drosophila blastoderm embryo.

**Figure 6:**
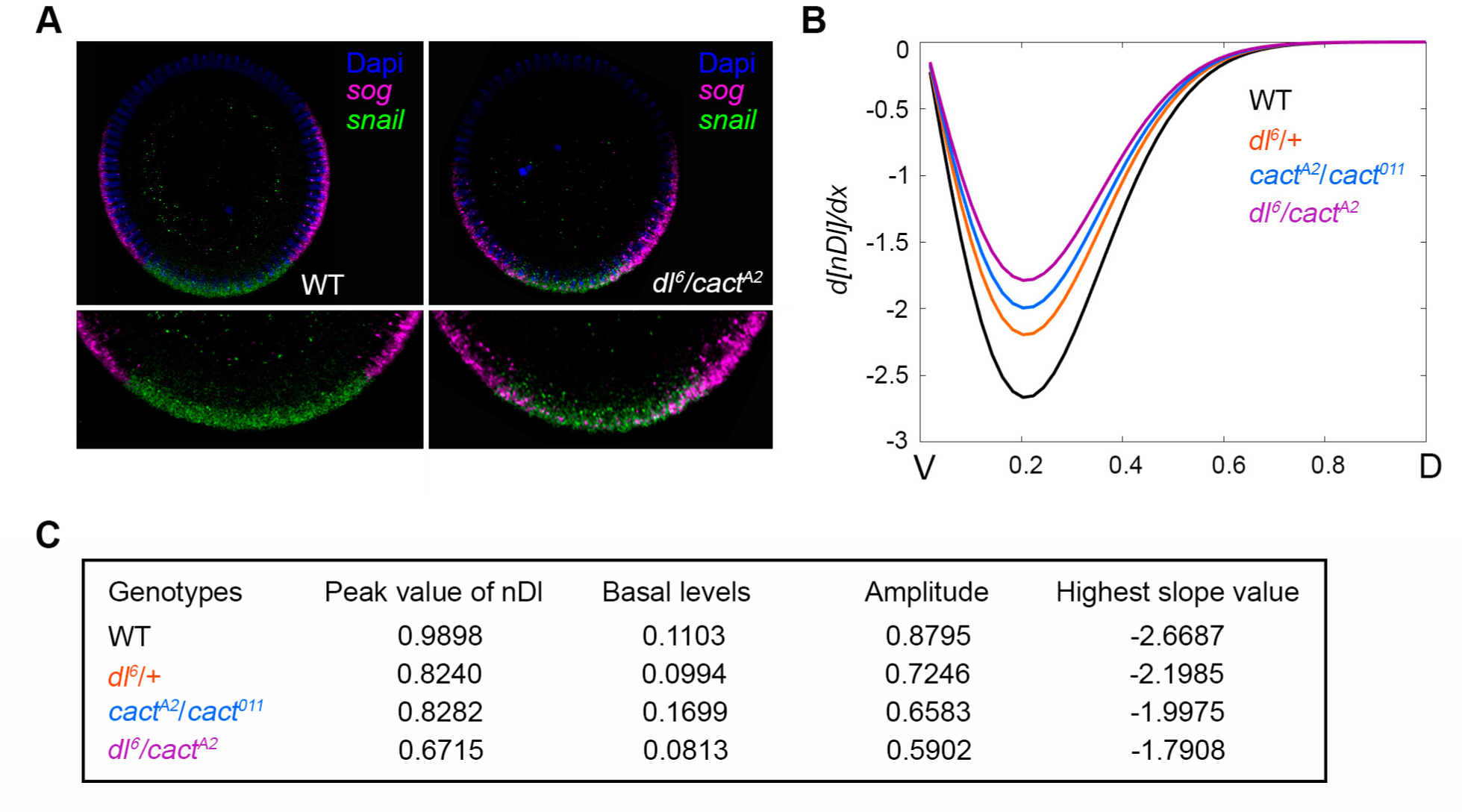
Variations in nDl gradient slope disrupts the precision of target gene expression DV territories. (A) Cross-sections of wild type (WT) and mutant (*dl^6^/cact^A2^*) cleavage cycle 14 embryos hybridized with *snail* (green) and *sog* (magenta) antisense RNA probes. Nuclei were stained using DAPI (blue). Note *sog* transcripts invading the ventral territory in *dl^6^/cact^A2^* mutants (zoomed images). (B) Derivatives of nDl concentration (y axis) as a function of the position along the embryonic DV axis (x axis) for WT (black), *dl^6^/*+ (orange), *dl^6^/cact^A2^* (purple), and *cact^A2^*/*cact^011^* (blue) mutant embryos. (C) Peak, basal levels, amplitude and highest slope of nDl distribution for each different genetic background.

### Model simulations reproduce Toll-dependent and Toll independent pathway mutant conditions

To challenge the predictive strength of our model we simulated different mutant conditions described in the literature and their effects on the nDl gradient. First, we simulated a *pelle* mutant (*pll-*). Pelle is a DEAD-domain kinase activated downstream of the Toll receptor, likely involved in Cact phosphorylation with consequent ubiquitination and degradation by the proteasome (Daigneault *et al*., 2013; Liu *et al*., 1997). Loss-of-function maternal *pll* mutants lead to dorsalized embryos, with loss of lateral and ventral elements of the cuticle (Galindo *et al*., 1995; Shelton and Wasserman, 1993). By reducing k9 or k10, the constants controlling Cact phosphorylation and release of Dl dimers for nuclear translocation (Fig. 1), nDl levels gradually decrease. At k9 or k10 = 0, all nuclei display the same level of nDl, equivalent to levels observed in the dorsal region of wild-type embryos (Fig 7A). Accordingly, the amount of DlCT and DlC accumulates (Fig. 7B,C), since Dl dimers are not released from Cact inhibition. This effect is restricted to ventral and lateral regions where pre-signaling complexes are recruited to active Toll. No effect is seen in the dorsal region where nDl results solely from direct Dl flow.

**Figure 7:**
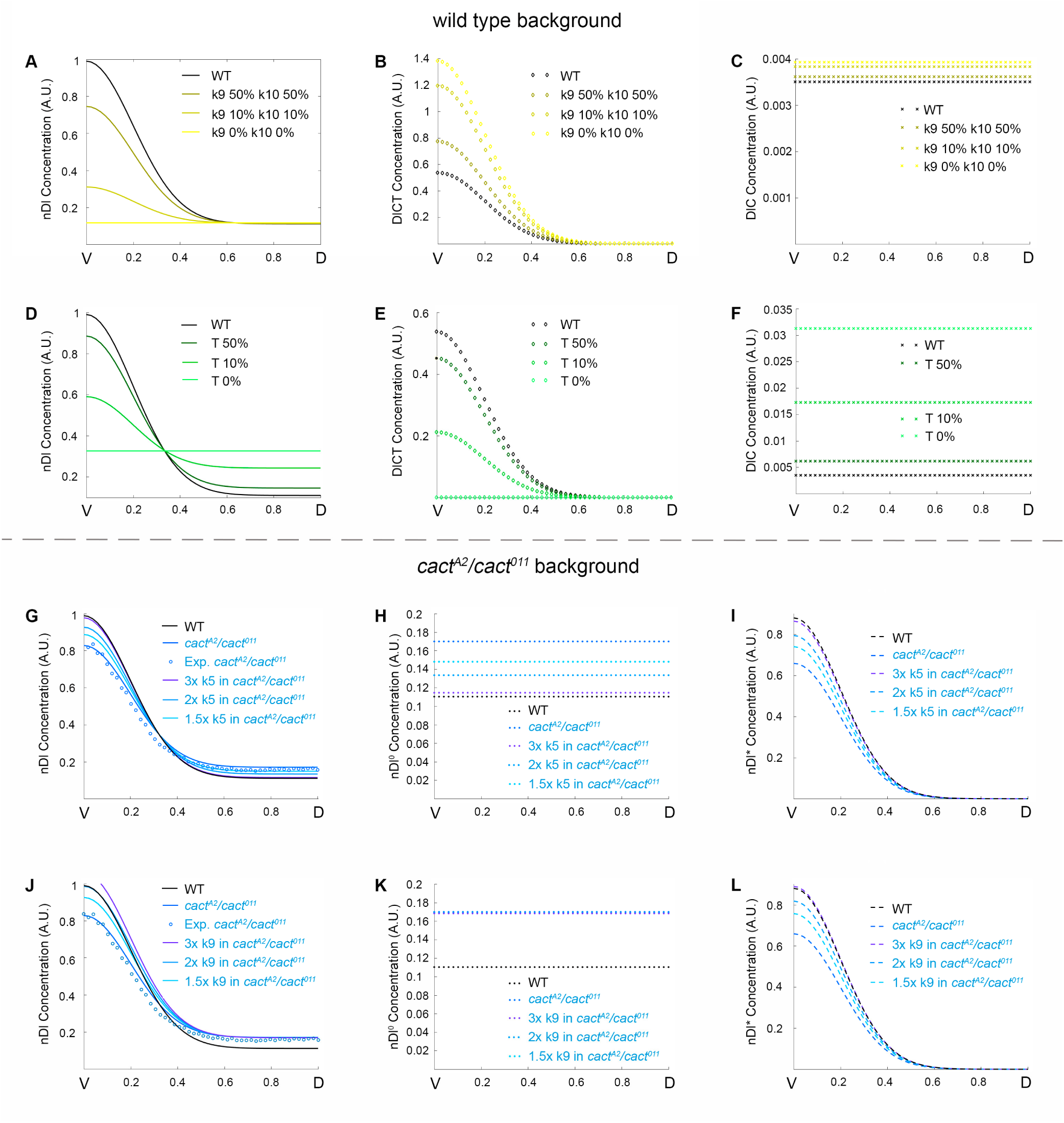
Model simulations reproduce Toll-dependent and Toll independent pathway mutant conditions. (A-C) Simulations for simultaneous reduction of kinetic constants k9 and k10 (0%, 10% and 50% reductions), showing the distribution of nuclear Dorsal (A), DlCT complex (B) and DlC complex (C). (D-F) Simulations decreasing Toll receptor (T) in 0%, 10% or 50%, showing the distribution of nuclear Dorsal (D), DlCT complex (E) and DlC complex (F). (G-L) Simulations increasing kinetic constant k9 (G-I) or k5 (J-L), by 3x, 2x or 1.5x, in a *cact^A2^*/*cact^011^* mutant background. Shown are nDl (total nuclear Dorsal dimers, G,J), nDl^0^ (free nuclear Dorsal dimers; H, K) and nDl* (nuclear Dorsal dimers activated by Toll Pathway; I, L). G and J also display experimental data from the *cact^A2^*/*cact^011^* mutant (circles) for comparison with wild type and mutant simulations. All concentrations are plotted along the V-D axis.

Different from *pll* loss-of-function, *Toll*[rm9] loss-of-function mutants lead to lateralized embryos, where the level of nDl is uniform along the entire DV axis and induces the expression of lateral Dl-target genes that require intermediate Dl levels for activation (Liberman *et al*., 2009). By simulating a decrease in the amount of activated Toll, we observe that the nDl gradient flattens to a uniform level along the DV axis (Fig. 7D). In addition, DlCT reduces while DlC increases progressively compared to wild type (Fig. 7E-F). Thus, the main difference we observe by simulating *Toll-* and *pll-* is that in *pll-* there is an increase in DlCT, while in *Toll-* the concentration of this complex reduces (Fig. 7E,F).

To explore the effects of alterations in the Toll-independent pathway for Cact regulation we changed kinetic parameters that are not directly related to Toll receptor activation, namely, k2 and k5. Decreasing any of these constants led to contrasting effects on dorsal versus ventral regions of the embryo, replicating the dual effect we observe in *cact*[A2]/*cact*[011] (Fig. S5E-H). Conversely, by increasing k5 we were able to recover the *cact*[A2]/*cact*[011] dual phenotype (Fig. 7G-I). A similar effect is observed by increasing k2 or k7 in the same *cact*[A2]/*cact*[011] background (Fig. S5 A-D). Recovery to the wild-type pattern is paralleled by a homogeneous decrease in nDl^0^ (Fig. 7H and Fig. S5B) and ventral increase of nDl* (Fig. 7I and Fig. S5C). Interestingly, this effect closely resembles that seen by providing excess Cact[E10]eGFP to *cact*[A2]/*cact*[011] embryos (Cardoso *et al*., 2017). This suggests that the N-terminal deleted Cact[E10] fragment inhibits basal nuclear translocation of Dl (nDl^0^) and favors nDl* by an unknown mechanism that should be explored in the future. We also simulated the effect of enhancing Toll signals in a *cact*[A2]/*cact*[011] background, by increasing k9. As a result, in ventral regions of the embryo nDl recovers the simulated *cact*[A2]/*cact*[011] gradients to wild-type levels. However, the gradient is unchanged in dorsal regions (Fig. 7J-L). Altogether, these results indicate the dual effect observed in *cact* mutants requires Cactus regulation by Toll-independent events.

## DISCUSSION

### Cactus plays a positive role for Dl nuclear transport by controlling the levels of Toll-responsive Dorsal-Cactus complexes

The *Drosophila* embryo is a unique setting for investigating Toll pathway function, as each genetically identical nucleus receives a unique level of Toll activation, distinct from its neighbor nuclei. Dorsal-ventral patterning of the embryo depends on this graded activity, instigating scientists to model the steps from Toll activation to formation of the nuclear Dorsal gradient. The predictive network model herein proposed adds novel elements to the regulatory network modeling of nDl gradient formation in the *Drosophila* embryo and expands our understanding of mechanisms that control NFkB/c-Rel activity.

First, the model enables us to explore the contribution of Toll-activated versus direct Dl nuclear translocation to the nDl gradient. In simulating a wild-type context, the model predicts a uniform level of nDl imported by the direct pathway, consistent with the previously reported uniform flow of dimers from cytoplasm to nucleus along the DV axis (DeLotto *et al*., 2007). According to model parameters, Dl nuclear translocation by the Toll pathway is more efficient than by direct flow, which is also supported by experimental data (Drier *et al*., 1999). Furthermore, by predicting the individual concentrations of all chemical species, the model indicates that the majority of cytoplasmic Dl is bound to either Cact and/or Toll receptor (DlC and DlCT), conforming to the relative levels of free versus Cact-bound Dl protein in embryonic extracts (Isoda and Nusslein-Volhard, 1994).

Second, the proposed model reproduces the dual effect of *cact* loss-of-function combinations and points out the essential components of the network that enable this dual effect. Broadly accepted as an NFκB family protein inhibitor with the role of securing Dl in the cytoplasm, our finding of a Cact function to favor Toll signals implies the existence of unexplored mechanisms that control Dl activity. Our simulations implicate the translational control of Cact levels by Dl protein and two modes of Dl nuclear translocation, Toll dependent and direct Dl nuclear translocation, as key aspects of Dl gradient formation. The most important components involved in these two routes are DlC and DlCT complexes. In forming DlC, Cact plays a negative role to impair Dl from entering the nucleus, while by enabling new DlCT complexes it favors Dl nuclear translocation. Since Toll activity is high ventrally, DlCT predominates in this region. Contrarily, in the dorsal region only DlC is observed. Therefore, simulations suggest that this differential distribution, together with the opposite role that Cact plays in each DV region, are the characteristics that enable Cact dual effect on Dl nuclear translocation (Fig. 4). The generation of endogenous *dl* and *cact* alleles that either increase or decrease the rate of association between Dl, Cact and Toll pathway adaptor proteins should enable to test these predictions by altering the relative amount of free protein versus protein complexes.

By simulating additional mutant conditions that decrease Cact function, our model confirms that the two modes of Dl nuclear translocation are the basis of the dual effect of Cact to favor Dl nuclear translocation ventrally and inhibit Dl nuclear translocation dorsally. These simulations indicate that the lower the expression of Cact, the greater the amount of nDl that enters the nucleus by direct flow, contributing to increase the basal (uniform) component of nDl concentration. Notably, increasing Toll pathway activity (increasing k9) does not recover the full *cact* loss-of-function phenotype. Only by modifying the constants associated to Toll-independent processes we were able to recover the phenotype to a wild-type pattern. This indicates that the Toll-independent pathway is fundamental for Cact to perform its different effects. It will be interesting to investigate in greater detail how the Toll-independent pathway controls basal Dl nuclear translocation and how it contributes to the nDl gradient.

### Two routes for Dorsal nuclear translocation as an extension of previous models

The pioneer model proposed by Kanodia et al. (2009) to explain Dorsal gradient dynamics during *Drosophila* embryogenesis was very successful in pinpointing the most important features of gradient formation during nuclear division cycles 11-14. This model also proved extremely powerful when comparing nDl gradient formation in related *drosophilids*. (Ambrosi *et al*., 2014). In this publication, Ambrosi also suggest that Cactus is the most important element that allows the correct gradient accommodation in embryos of greatly divergent sizes. However, previous models confer Cactus only an inhibitory role, where less Cactus in the cytoplasm is expected to decrease Cactus-Dorsal association, favoring Toll dependent Dorsal nuclear localization. Inspired by the fact that in certain Cactus loss-of-function mutants nuclear Dorsal concentration decreases in ventral regions, we have proposed a two-step model that accounts for this peculiarity. First, the concentration of Cactus-Dorsal complex is controlled by a Toll-independent mechanism involving Cactus synthesis and processing or recruitment into a complex with Dl. Second, the DlC complex associates with Toll, which promotes facilitated entry of Dorsal into nuclei and irreversible Toll-dependent Cactus degradation. In this case less Cactus results in less DlC + DlCT complexes and consequently, contrary to the previous model, less Toll-induced Dorsal entry into the nuclei whenever the balance between Toll signals and availability of new DlC + DlCT signaling complexes is perturbed. The general result of this model is that nuclear Dorsal follows the Toll activation gradient plus a basal uniform concentration due to the direct entry of Dorsal independent of Toll, also introduced in the model.

A fundamental feature of our network that enables to simulate Cactus’ positive function is that the formation of DlC complexes depends not only on Toll activation but also on association-dissociation events that may be controlled by Toll-independent inputs. Variation in the amount of DlC complexes may explain why the free flow of Dorsal into the nucleus (nDl^0^) is almost unaffected in *dl[6]/+* mutants, compared to the significant decrease in Toll-induced nDl (nDl*). DlC decreases roughly 30% as a result of a 15% decrease in total Dl protein (Fig. 3C, inset), whereas free Dl reduces only 10% (cDl^0^ and nDl^0^; Fig. S1) as there is less Cact to form DlC. On the other hand, in the ventral side of the embryo 20% less nDl enters the nucleus in response to Toll activation (nDl*), since there is less DlC to form new Toll responsive complexes (DlCT). Accordingly, in the ventral domain of *cact*[A2]/*cact*[011] embryos, DlC decreases 50%, leading to a great decrease in DlCT and Toll-induced Dl nuclear translocation, although direct Dl flow increases (Fig. S2 and Fig. S4). It is noteworthy that, since the ratio between kinetic constants for Toll-dependent Dl nuclear transport (k11/k12 = 545.45) is higher than via direct flow (k3/k4 = 1.95), the Toll induced Dl nuclear translocation process is more efficient than Dl free flow. Therefore, alterations in DlC affect differently the two Dl nuclear translocation routes.

The observation that DlC is central to explaining the behavior of the different mutant conditions hereby analyzed points to the importance of identifying regulators of DlC formation. Processes that control the mobilization of DlC to mount new Toll-signaling complexes (DlCT) or that alter the availability of Dl and Cact for association-dissociation events are likely to induce unanticipated effects on NFκB family transcription factor activity. Among the signals that control Cactus independent of Toll, Calpain A, that impacts the nDl gradient (Fontenele *et al*., 2013), may also control the formation of DlC and DlCT complexes since it localizes close to the membrane where these events should take place (Fontenele *et al*., 2009). Future investigations on the effect of Calpain A and other Toll-independent pathway regulators on DlC levels and free Dl and Cact may shed light on these important issues.

Further extensions of the Kanodia model were proposed in the literature by taking into account the presence of Cactus in the nuclei (O’Connell and Reeves, 2015) and by implying diffusion rates that lead to shuttling of Cactus-Dorsal along the DV axis (Carrell *et al*., 2017). We show that our model simulations correctly reproduce nDl properties displayed in several mutants, without need of any additional assumption.

### The balance between reaction and diffusion of free and complexed Dorsal and Cactus

Molecular diffusion of proteins involved in the response to Toll signals greatly impacts the nuclear Dorsal concentration gradient. Our model suggests that, due to interaction with the Toll receptor, the DlC complex diffusivity is lower than free Dl and Cact (Table 1). Paradoxically, cDl^0^, nDl^0^, Cact and DlC show uniform distribution along the embryonic DV axis (Fig. S1A-D), despite interacting with non-uniformly distributed DlCT and nDl*. The answer to this paradox lies in the balance between lateral (DV) diffusion and Dl nuclear translocation. It is important to note that both cDl* and cDl^0^ translocate to the nucleus, an action that competes with movement to the adjacent lateral compartment. However, the intercompartmental diffusion term D_Dl_ / L^2^ (L = 1/50 on the normalized scale, hence D_Dl_ / L^2^ ≈ 227.500 in inverse time units) is much larger than the kinetic constants k3, k5 and k11 (direct flow, DlC complex formation and Toll mediated Dl nuclear entrance, respectively). Thus, diffusive dynamics throughout the cytoplasm dominates the kinetics of cDl^0^, leading to uniform free Dorsal throughout the embryo, even though total nuclear Dl obeys a gradient. We found a similar scenario for Cactus, given that D_C_ / L^2^ ≈ 1.037.500 (in inverse time units) is much larger than kinetic constant k2 and k5. We found an opposite behavior for the Dorsal-Cact complex, which is not as diffusible as the free forms. It is noticeable that D_DlC_ / L^2^ ≈ 0.08475 is much lower than k11, which favors the entry of Dl into the nucleus via the Toll pathway in the ventral and lateral regions. This value is also much lower than k6, the reversible process that returns the complexed form DlC to the free forms cDl^0^ and C_f_. Therefore, low DlC diffusivity favors the kinetics of DlC binding to the Toll receptor, making the DlCT gradient strongly correlated to the Toll activation profile.

It has been previously reported that the diffusion between adjacent compartments takes place much more slowly than diffusion within individual nucleo-cytoplasmic compartments during mitosis (Daniels *et al*., 2012), since the cytoplasm surrounding each nucleus is partially compartmentalized (DeLotto *et al*., 2007). Also, Kanodia and collaborators (Kanodia *et al*., 2009) assumed that the diffusion coefficient of Dorsal-Cact complex (DlC), free Dorsal and Cact are identical, resulting in a lateral transport coefficient of approximately one. This indicates that each of the molecular species above traverses a single compartment during cycle 14. Besides that, even reporting different diffusion coefficients for Dorsal and Cactus, (Carrell *et al*., 2017) estimate that Dorsal travels 7-10 compartments over a 90-minute period (cycles 10 to 14), and then argues that, taking into account the changing distances between nucleo-cytoplasmic compartments due to mitosis, the transport coefficient is centered around 1. We found a similar result for the lateral transport coefficient for the DlC complex (λ = D_DlC_ T / L^2^ ≈ 5 where T = 60 minutes is the total cycle 14 time duration). However, this value may be overestimated since we simulated only the 14th mitotic cycle, setting a uniform distribution of cytoplasmatic free Dorsal cDl^0^ as the initial condition (see Methods for more details). Furthermore, along cycle 14 membrane furrows extend and close around each nucleus, limiting further the extent of lateral diffusion.

The above results indicate that our model simulations agree with the confinement of Dorsal within individual nucleo-cytoplasmic compartments as proposed by others, but indicates that this holds true only for the Dorsal-Cact complex. The diffusion of individual species Dl and Cact, on the other hand, plays a critical role for the balance between reaction and diffusion required to correctly establish and maintain the Dl nuclear gradient, as described above.

### A conserved regulatory network for Cactus/IκB function

Toll was initially described in *Drosophila* embryos as a maternal effect allele regulating DV patterning (Anderson *et al*., 1985b). However, it has been suggested that the function of Toll to pattern the DV axis is restricted to the insect lineage, while regulating the innate immune response is an ancient and widespread function (Benton *et al*., 2016; Lynch and Roth, 2011). In either context, elements of the Toll pathway are conserved in Bilateria. Conservation is not restricted to elements downstream of the Toll receptor. Toll-dependent and -independent pathways have been reported to regulate vertebrate IκB, as shown for *Drosophila* Cact. In mammalian cells, it has been proposed that constitutive degradation of IκBα independent of the proteasome regulates basal levels of IκB and thus the duration of the NFκB response (Han *et al*., 1999; Karin and Ben-Neriah, 2000; Shen *et al*., 2001; Shumway *et al*., 1999). Experimental data and modeling of signal independent IκB regulation suggest that the regulatory pathway controlling free IκB is a major determinant of constitutive NFκB and of stimulus responsiveness of the NFκB signaling module (O’Dea *et al*., 2007). Furthermore, calcium-dependent Calpain proteases also target mammalian IκB independent of Toll receptors (Schaecher *et al*., 2004; Shen *et al*., 2001; Shumway *et al*., 1999), as reported for Cact in the embryo and immune system (Fontenele *et al*., 2009; Fontenele *et al*., 2013). Unlike *Drosophila*, vertebrates rely on several IκB proteins, where the final effect on the immune response over time is a result of the compounded effect of three or more IκBs (Kearns *et al*., 2006). Interestingly, it has been shown that IκBβ knockout mice display a dramatic reduction of TNFα in response to lipopolysaccharide (LPS), suggesting that IκBβ acts to both inhibit and activate gene expression (Rao *et al*., 2010). Even though the molecular mechanism proposed by Rao et al for IκBβ is different from the here presented for Cact, they are similar in the sense that both are able to improve NFκB family transcriptional activity. Therefore, there is an open avenue of investigation on the mechanisms that regulate Cact/IκB function with potentially important outcomes for immunity. Altogether, the results herein presented credit to IκB/Cact protein a key role in NFκB activity greater than previously reported.

## METHODS

### Mathematical modeling of Dl nuclear gradient formation

The proposed reaction-diffusion model is a refinement of (Kanodia *et al*., 2009), in which the authors described Dl and Cact dynamics by a reaction-diffusion regulatory network along a 1D spatial domain that represents embryo’s outline in a dorsal-to-ventral cross section. The pioneer model was used and modified by others (Carrell *et al*., 2017; O’Connell and Reeves, 2015). Here we expanded this model to take into account the translational control of Cact levels by Dl protein and the two different routes for Dorsal nuclear localization.

We assumed that the embryo is symmetric with respect to the DV axis, therefore, all simulations acknowledge only one of the embryo’s dorsal-ventral cross-section half, considering no-exchange boundary conditions at both ventral and dorsal midlines. The number of compartments is fixed and their sizes are equal, such that the length of the whole simulation region is one, for computational convenience.

Dorsal dimerization was not described in the model, since nuclear input dynamics is restricted to the dimeric form. Thus, although the model refers simply to Dl, it should be kept in mind that, for the purposes of this work, the species in question is always the corresponding dimer. The association of cytoplasmic Dorsal dimer (cDl°) with Cactus (C_f_) is also described and the resulting complex is treated as a new species (DlC). In addition, direct nuclear Dorsal import and export is treated as a reversible reaction in which nuclear Dorsal is treated as a new nuclear species (nDl°).

The Toll receptor (T) in the model is the one activated by Spätzle, and hence its distribution along the dorsoventral region in the embryo is not uniform. In this work, it was assumed that its distribution is Gaussian, whose mean value is in the most ventral part of the embryo, and its standard deviation and intensity peak are two parameters fitted to experimental data, along with the kinetic constants and the diffusion coefficients.

In addition to participating in the reactions, Fig. 1, some of the proteins can diffuse along the compartments. In particular, cDl°, C_f_ and DlC can diffuse following Fick’s Law whereas cDl* and C_ub_ reaction dynamics are considered much faster than their diffusion capacities, hence they do not have diffusion coefficients associated with them. All of the processes, which are modelled as chemical reactions, take place inside each compartment and are described by the Mass Action Law. Further details about the resulting differential equations are displayed in Supplementary Text.

### Model calibration and simulation

To determine the model parameters, we calibrated kinetic parameters k1 to k12, cDl°, C_f_ and DlC diffusion coefficients, total Dl concentration, and Toll receptor intensity peak and standard deviation (assuming that its activation follows a Gaussian function centered at the most ventral region of the embryo) using experimental data of nuclear mitotic cycle 14 wild-type *Drosophila* embryos (Cardoso *et al*., 2017). We set up an optimization problem such that the objective function is the quadratic loss where the model total nuclear Dl (the sum of nDl^0^ and nDl*) was compared to experimental nuclear Dorsal data. The sum of the quadratic differences between simulation and experimental data was minimized. The experimental data used was normalized so that the peak of nDl concentration at the most ventral midline is 1.

Since the loss function is not explicitly defined in terms of the parameter because it results from the numerical solution of a Partial Differential Equations System (PDEs), metaheuristics become convenient for intelligently exploring parameter possibilities, saving computational cost to obtain a viable set of parameters. Among several techniques, we have adopted the Genetic Algorithm (GA), which was implemented using a binary codification of the parameters, a selection by a k-tournament, a uniform bit-wise crossover and a mutation by bit-wise inversion (Gen and Lin, 2008; Mitchell, 1998).

For each individual in the Genetic Algorithm population, which corresponds to a set of parameters to be evaluated and ranked by the objective function, we have solved the PDEs by doing a second-order central approximation to deal with spatial coordinates, giving rise to an ODE system for each compartment. Subsequently, a linear implicit multistep method based on 6^th^ order finite differences (BDF-6 formula) was used to solve each ODE in time (Ashino *et al*., 2000; Shampine and Reichelt, 1997). The computation was done such that it ended when the system had reached a steady state for each individual in the GA population.

After running the program and finding out an optimized set of parameters, the mutant simulations have been done by changing specific parameters and comparing the newer simulations with determined set of experimental data (Cardoso *et al*., 2017). For *dl6/+* mutants, we have only changed total Dl concentration; for *cact-*related mutants, kinetic constant k1, which is related to Dl regulation of Cact, was changed; and both of these parameters were changed for *dl^6^/cact^A2^* mutant. Further details about the equations, GA’s parameters and other specifications can be found in Supplementary Text.

## Acknowledgments

We thank Trudi Schupbach for helpful comments on the manuscript.

## Funding

FL and HA was supported by FAPERJ - Fundação de Amparo à Pesquisa do Estado do Rio de Janeiro [Grant no. E-26 010.001877/2015].

