## Supplementary Text for "A Reaction-Diffusion Network model predicts a dual role of Cactus/IκB to regulate Dorsal/NFκB nuclear translocation in *Drosophila*"

**Model equations and parameters.** Fig. 1A illustrates the proposed reaction-diffusion model, in which transport from one compartment to another is given by Fick's Law, and reactions are given by Mass Action Law. Nuclear import and export are modeled as reactions that convert cytoplasmatic Dl to nuclear Dl. Total Dl concentration was a calibration parameter assumed to be uniform along the embryo; activated Toll was assumed to exhibit a Gaussian shape function with two parameters A and σ:

$$[T_{tot}] = A \cdot exp\left(-\frac{x^2}{\sigma^2}\right) \quad (1)$$

The partial differential equations system (PDEs) governing that system's dynamic is given by:

$$\frac{\partial[cDl°]}{\partial t} = D_{cDl°}\frac{\partial^2[cDl°]}{\partial x^2} - k_3[cDl°] + k_4[nDl°] - k_5[cDl°][C_f] + k_6[DlC] + k_{12}[nDl*] \quad (2)$$

$$\frac{\partial[C_f]}{\partial t} = D_{C_f}\frac{\partial^2[C_f]}{\partial x^2} + k_1[cDl°] - k_2[C_f] - k_5[cDl°][C_f] + k_6[DlC] \quad (3)$$

$$\frac{\partial[DlC]}{\partial t} = D_{DlC}\frac{\partial^2[DlC]}{\partial x^2} + k_5[cDl°][C_f] - k_6[DlC] - k_7[DlC][T] + k_8[DlCT] \quad (4)$$

$$\frac{\partial[T]}{\partial t} = -k_7[DlC][T] + k_8[DlCT] + k_9[DlCT] \quad (5)$$

$$\frac{\partial[DlCT]}{\partial t} = k_7[DlC][T] - k_8[DlCT] - k_9[DlCT] \quad (6)$$

$$\frac{\partial[C_{ub}]}{\partial t} = k_9[DlCT] - k_{10}[C_{ub}] \quad (7)$$

$$\frac{\partial[cDl*]}{\partial t} = k_9[DlCT] - k_{11}[cDl*] \quad (8)$$

$$\frac{\partial[nDl°]}{\partial t} = k_3[cDl°] - k_4[nDl°] \quad (9)$$

$$\frac{\partial[nDl*]}{\partial t} = k_{11}[cDl*] - k_{12}[nDl*] \quad (10)$$

**Loss Function:** In order to find the parameters that minimize the differences between experimental and simulated nuclear Dl, the following loss function was defined:

$$L = \frac{1}{n}\sum_{i=1}^{n}([nDl]_i^{exp} - ([nDl^{\circ}]_i + [nDl*]_i))^2 \quad (11)$$

where $n$ is the total number of compartments (in our case, $n = 50$) and $[nDl]_i^{exp}$ is the experimental data of total nuclear Dl in compartment $i$ (which ranges from 1 to the total number of compartments $n$).

The loss function implicitly depends on all parameters, since it depends on simulated nDlº and nDl* levels. Therefore, one must minimize the loss function to find a parameter set that approximates computational results to experimental data.

**Genetic Algorithm usage and parameters:** Each 12 kinetic constants, 3 diffusion coefficients, total Dl concentration and 2 activated-Toll related parameters have been coded by 12 bits. Each population consisted of 50 individuals, the Tournament Size used was k = 2, the recombination probability for each individual was 75%, and thus the exchange probability given cross over was 60% for each bit pair. In addition, the mutation rate per bit was 2%. The elitism percentage considered was 5%, which guarantees survival of 3 individuals per generation, and the algorithm run for 500 iterations.
