## Supplementary Figure 2 for "A Reaction-Diffusion Network model predicts a dual role of Cactus/IκB to regulate Dorsal/NFκB nuclear translocation in *Drosophila*"

**Barros et al. Supplementary Figure 2**

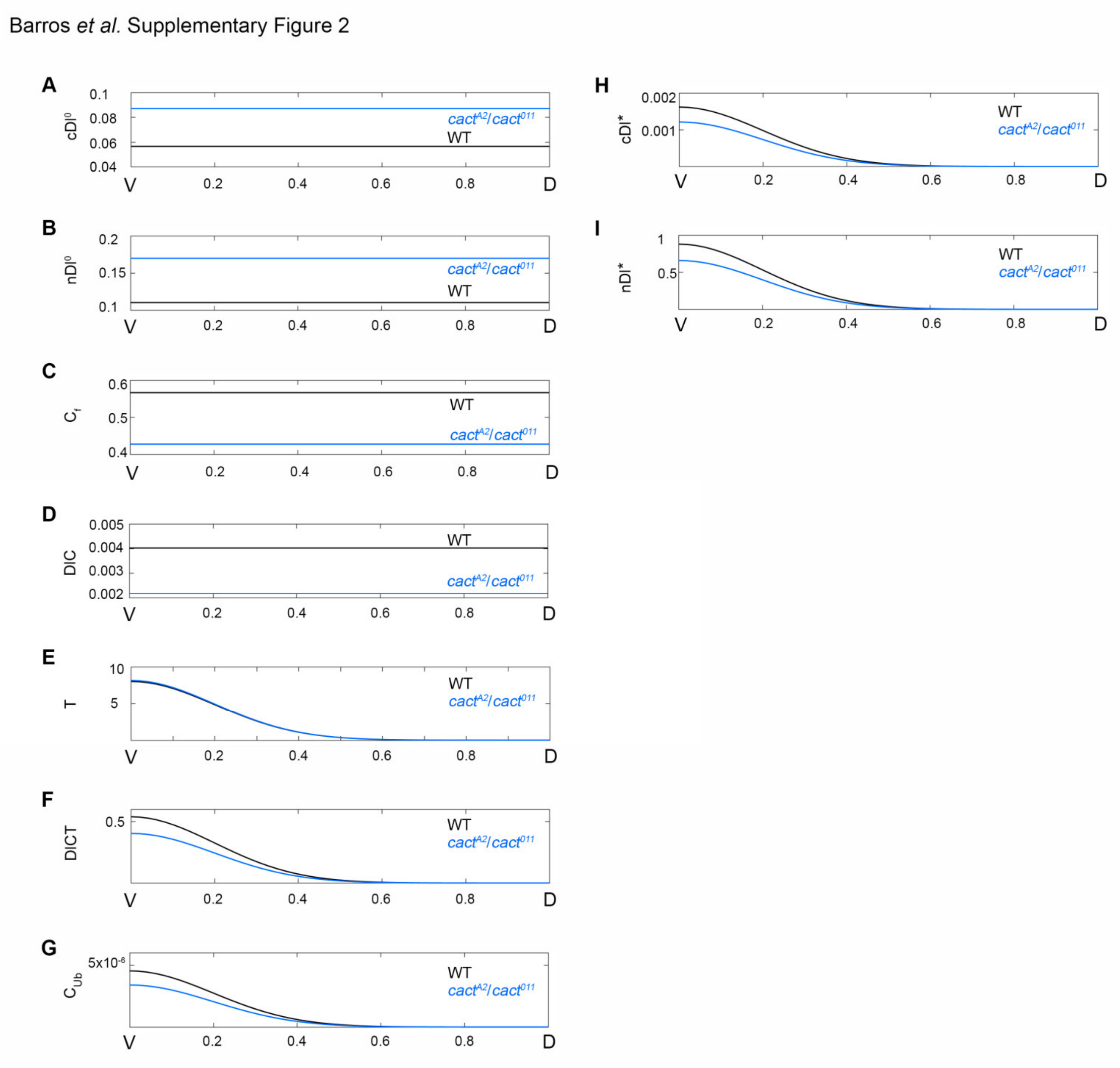


**Figure S2: Model prediction of dorso-ventral distribution of all model species for wild type and *cact^A2^/cact^011^* mutant.** Distribution of free cytoplasmic (cDl$^0$, A) and nuclear (nDl$^0$, B) Dorsal, free Cactus ($C_f$, C), DlC complexes formed by Dl dimer and Cact monomer (D), Toll (T) receptor (E), DlCT complexes including DlC and a Toll receptor (F), Cactus ($C_{ub}$, G) and cytoplasmic Dorsal (cDl*, H) modified by Toll Pathway (G-H), nDl modified by Toll induced (I).
