## Supplementary Figure 3 for "A Reaction-Diffusion Network model predicts a dual role of Cactus/IκB to regulate Dorsal/NFκB nuclear translocation in *Drosophila*"

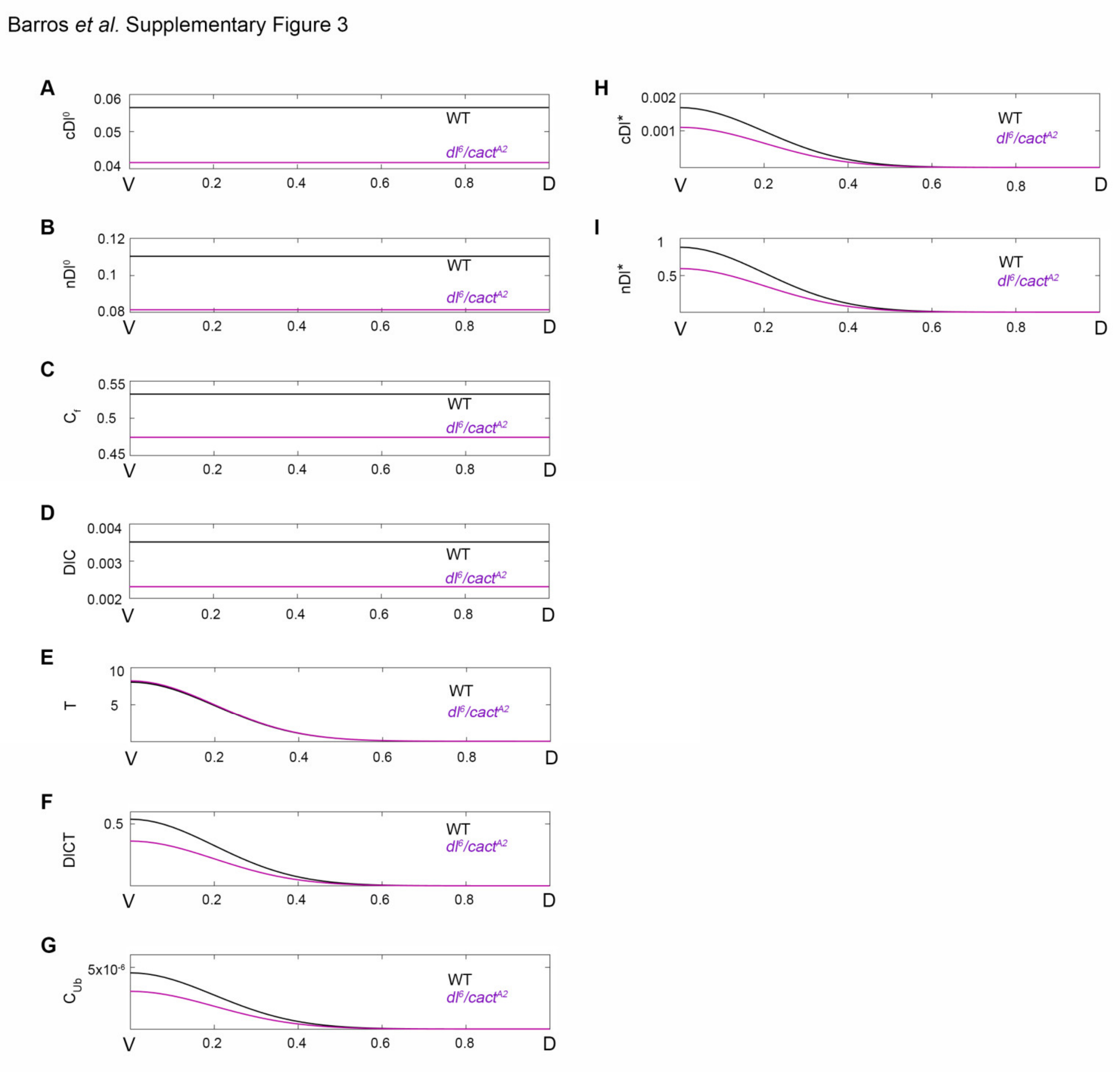


**Figure S3: Model prediction of dorso-ventral distribution of all model species for wild type and *dl⁶/cact*$^{A2}$ mutant**. Distribution of free cytoplasmic (cDl$^0$, A) and nuclear (nDl$^0$, B) Dorsal, free Cactus ($C_f$, C), DlC complexes formed by Dl dimer and Cact monomer (D), Toll (T) receptor (E), DlCT complexes including DlC and a Toll receptor (F), Cactus ($C_{ub}$, G) and cytoplasmic Dorsal (cDl*, H) modified by Toll Pathway (G-H), nDl modified by Toll induced (I).
