## Supplementary Figure 4 for "A Reaction-Diffusion Network model predicts a dual role of Cactus/IκB to regulate Dorsal/NFκB nuclear translocation in *Drosophila*"

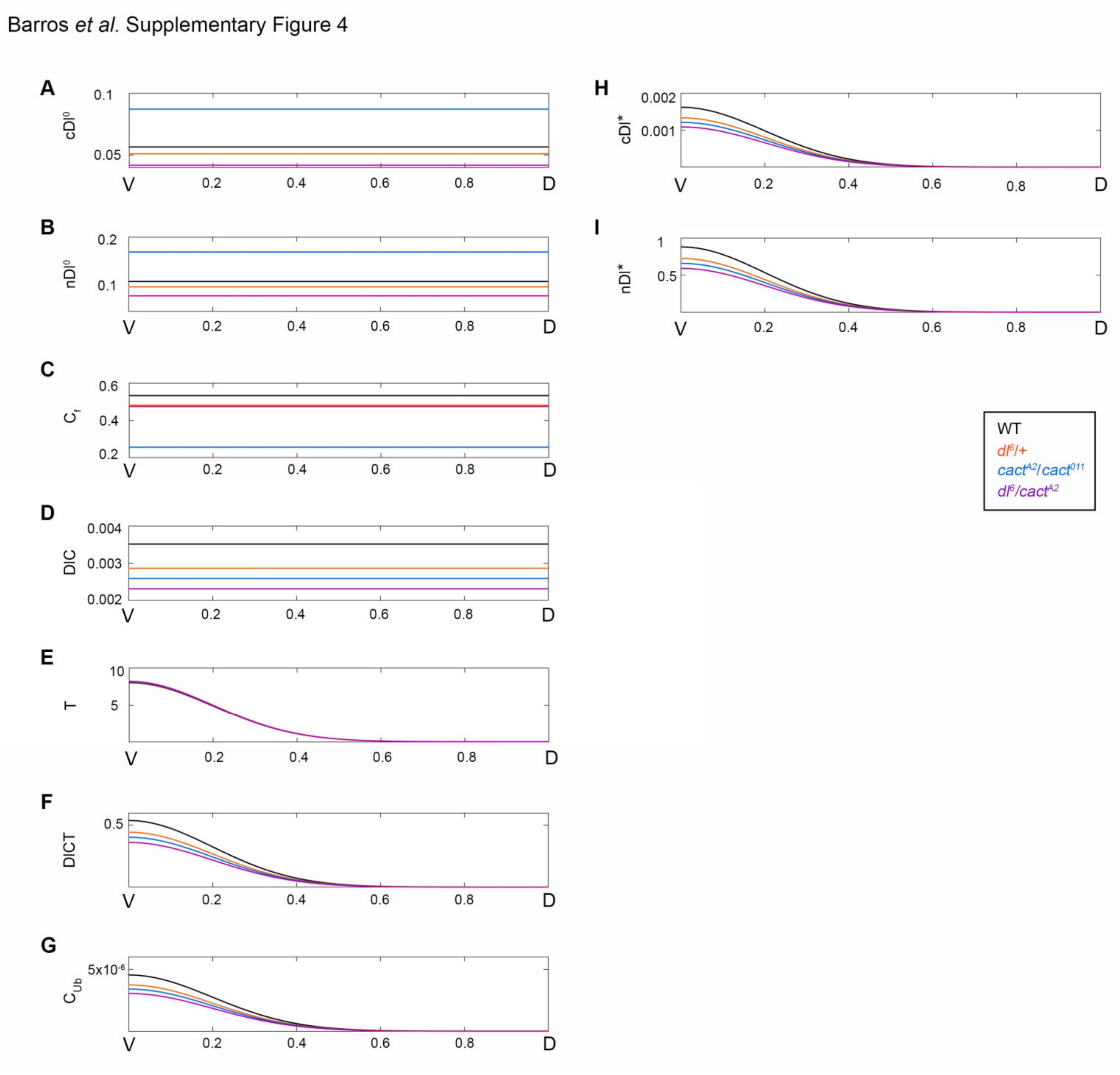


**Figure S4: Model prediction of dorsal-ventral distribution of all model species.** Curves for wild-type (black), *$dl^{6}$/+* (orange), *$cact^{A2}/cact^{011}$* (blue), and *$dl^{6}/cact^{A2}$* (purple) mutants. Distribution of free cytoplasmic ($cDl^{0}$, A) and nuclear ($nDl^{0}$, B) Dorsal, free Cactus ($C_{f}$, C), DIC complexes formed by Dl dimer and Cact monomer (D), Toll (T) receptor (E), DICT complexes including DIC and a Toll receptor (F), Cactus ($C_{ub}$, G) and cytoplasmic Dorsal (cDl*, H) modified by Toll Pathway (G-H), nDl modified by Toll induced (I).
