## Supplementary Figure 5 for "A Reaction-Diffusion Network model predicts a dual role of Cactus/IκB to regulate Dorsal/NFκB nuclear translocation in *Drosophila*"

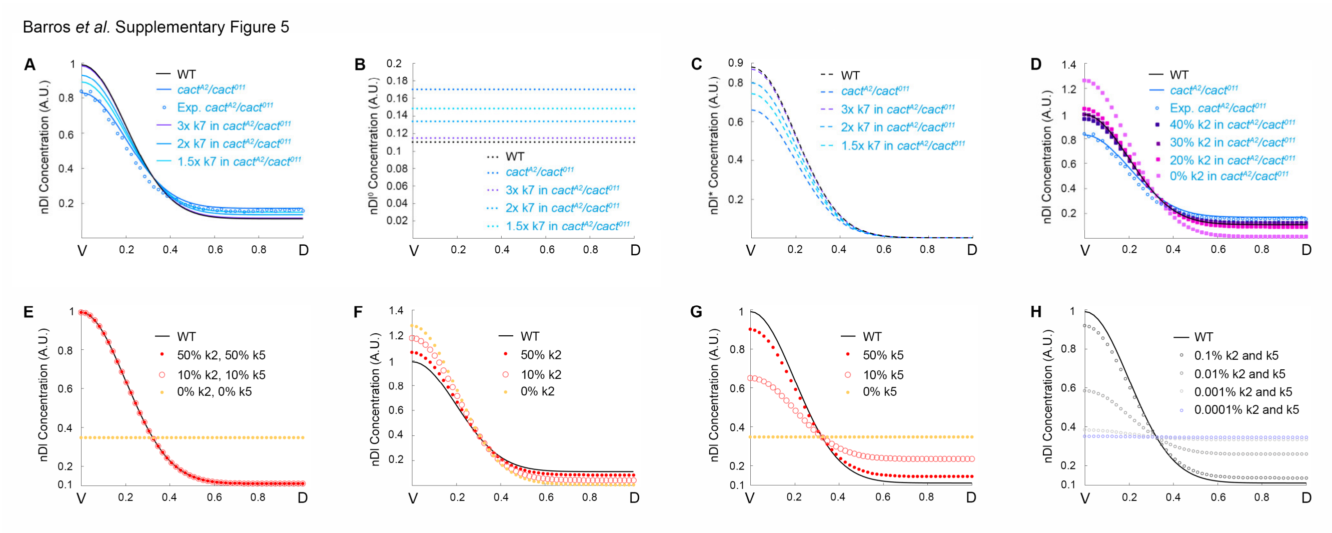


**Figure S5: Model simulations recover the dual effect of Cactus on Dorsal in *cact*$^{A2}$/*cact*$^{011}$ mutant background.** (A-C) Model prediction of nuclear species distribution for 3x, 2x or 1.5x increases of kinetic constant k7 in a *cact*$^{A2}$/*cact*$^{011}$ mutant background. nDl, total nuclear Dorsal dimers (A); nDl$^{0}$, free nuclear Dorsal dimers (B); nDl*, nuclear Dorsal dimers induced by Toll (C). (D) nDl simulation decreasing k2 by 20%, 30%, 40% or nulling k2 (0%) in a *cact*$^{A2}$/*cact*$^{011}$ mutant background comparing to experimental data (circle symbol) and WT and *cact*$^{A2}$/*cact*$^{011}$ mutant simulations. Simultaneous reduction of k2 and k5 by 0%, 10% and 50% (E) or by 0.1%, 0.01%, 0.001% and 0.0001% (H) comparing to control (WT) nDl concentration simulations (E, H). Reduction of k2 (F) or k5 (G) by 0%, 10% and 50%.
